# History of Traumatic Brain Injury with Loss of Consciousness and APOE ε4 Carriers Synergistically Increase Late-Life Amyloid PET Burden

**DOI:** 10.64898/2026.04.14.717801

**Authors:** Jeremy F. Strain, Nicolas R Barthélemy, Ruchira Jha, Owen Guo, Madhur Parihar, Kyle Chan, Babatunde Adeyemo, Peter R. Millar, Kyle Womack, Brian Gordon, Suzanne Schindler, John C. Morris, Tammie Benzinger, Beau M. Ances, Chia-Ling Phuah

## Abstract

**Background:** Traumatic brain injury with loss of consciousness (TBI-LOC) is an established risk factor for dementia, yet the pathways linking remote TBI to Alzheimer’s disease (AD) biology remain incompletely defined. APOE ε4 is the strongest genetic predictor of amyloid accumulation in late-onset AD, it may moderate the long-term consequences of head injury. This study investigates whether history TBI-LOC independently contributes or synergistically interacts with APOE ε4 to amplify late-life amyloid and tau burden.

**Methods:** 429 participants completed the Ohio State University TBI screening tool and an amyloid PET scan (centiloids). A subcohort (n=352) also underwent tau PET. TBI history was classified by recency (<10 vs >10 years) and severity (no TBI, dazing/confusion [TBI-DZ], TBI-LOC). Analyses were stratified by degree of clinical impairment as assessed by Clinical Dementia Rating (CDR=0 vs CDR>0). Logistic and linear regression models examined associations between TBI and amyloid, adjusting for age, sex, education, and APOE ε4, including an APOE*TBI-LOC status interaction term, while Fisher’s exact tests evaluated TBI recency and biomarker positivity.

**Results:** In CDR=0 participants (n=365), 119 reported a history of TBI, comprising 56 TBI-DZ and 63 TBI-LOC. TBI-LOC but not TBI-DZ, correlated with elevated amyloid PET levels (p<0.001; [4.6-17]). Furthermore, an interaction between APOE ε4 and TBI-LOC indicated that TBI-LOC augmented the amyloid-related risk associated with the APOE ε4 allele (p=0.003; [4.3-21]). The interaction persisted when stratified by TBI recency with only remote TBI-LOC (occurring more than 10 years prior) associated with increased amyloid PET (p=0.003 [5.2-25]). No association between TBI and tau was identified in a subset with tau PET, and no TBI-amyloid correlations were observed among symptomatic participants (CDR>0; n=64) suggesting a ceiling effect of pathology once clinical dementia is present.

**Conclusions:** History of remote TBI-LOC is linked to elevated amyloid PET levels in later life, particularly among APOE ε4 carriers with a CDR=0. The robust findings for amyloid, contrasted with null tau results and the reduced association in symptomatic cases underscore the importance of considering TBI history when screening for preclinical AD and assessing early-stage risk.

## Introduction

Traumatic brain injury (TBI) has been linked to a heightened risk of dementia in later stages of life[1]. Beyond its immediate effects, an increasing body of evidence indicates that a single TBI event may trigger or enhance mechanisms that contribute to pathological levels of amyloid accumulation later in life—a defining characteristic of Alzheimer disease (AD)[2]. AD is principally characterized by the progressive accumulation of pathology, initiated with amyloid-β plaques subsequently leading to the formation of neurofibrillary tangles (NFT). This well-established biomarker framework is often used as a benchmark to not only differentiate from other forms of dementia but to monitor progression. Pathological levels of amyloid have traditionally been used to define a preclinical stage and frequently as a criterion for clinical trials, with prior TBI history considered as a modifier for age of onset rather than a direct mechanism for amyloid accumulation. Evidence from pathological studies indicate the presence of amyloid deposition in acute TBI [3], but whether these pathological changes persist with chronically is less clear [4], [5]. Although this pathological change is present, amyloid associated with TBI is observed in roughly 30% of moderate to severe TBI cases and differentiating individuals at risk of developing amyloid accumulation later in life presents significant challenges[3], [6]. Moreover, TBIs exhibit significant heterogeneity, with secondary factors (TBI severity, genetic predisposition for AD, etc) likely modifying risk of AB deposition [7].

TBI is commonly considered as a broad categorization to encompass various conditions that arise as a result of head injuries. These conditions include cognitive state changes that can range in severity from mild confusion or disorientation to complete loss of consciousness and are assessed through self-reported accounts when medical records are unavailable[8]. Previous research has demonstrated that the severity of TBI may impact the risk of developing dementia[9]. Severe TBI has the potential to trigger long-term neurodegenerative changes, ultimately resulting in AD-like pathologies[10]. Large epidemiological studies utilizing administrative outcome measures have demonstrated an accelerated onset of cognitive impairment and Alzheimer’s disease-like pathology following moderate TBI in comparison to mild TBI[11] [12]. The likelihood of both acute and chronic impairments following a TBI is significantly heightened by the severity of the injury, frequently accompanied by a loss of consciousness.

Increased amyloid levels serve as the primary biomarker for pre-clinical staging of AD. Postmortem analyses of long-term TBI survivors have revealed widespread increases in amyloid accumulation even decades after the injury. The principal evidence underpinning this relationship originates from pathology and animal research[6], [13], [14], [15]; however, in-vivo PET studies have yielded inconsistent finings[5], [16]. PET is regarded as the gold standard for identifying pathological levels of AD-associated biomarkers and understanding the inconsistencies with prior exposure to TBI will increase its relevance in future clinical trials. Studies that do not identify a relationship often rely on visual assessments to quantify amyloid severity[5] or lack further consideration of additional secondary influencing factors like APOE ε4[17]. Mixed-method approaches have demonstrated an interaction between the APOE ε4 genotype, the principal genetic risk factor for late-onset AD, and TBI. These findings indicate that individuals possessing the APOE ε4 risk allele exhibit an elevated risk of developing AD after experiencing TBI in comparison to non-carriers[18]. Building on the hypothesis that TBI may initiate a feed-forward loop contributing to the subsequent onset of amyloid accumulation[19], it can be postulated that individuals with a higher genetic predisposition to AD are at greater risk after experiencing TBI compared to those with a lower genetic vulnerability[20].

Emerging evidence in this field increasingly highlights that the severity and recency of TBI, combined with genetic predispositions, represent critical factors that may heighten the likelihood of developing pathological levels of amyloid burden. Each of these potential contributing factors were evaluated within an enriched cohort for AD stratified by clinical presentation, using extensive positron emission tomography (PET) imaging to assess amyloid and tau severity, APOE ε4genotype, and a documented history of TBI.

## Materials and methods

### Study Cohort

The Charles and Joanne Knight Alzheimer’s Disease Research Center at the Washington University School of Medicine in St. Louis (WUSM) facilitated the recruitment of participants. The inclusion criteria necessitate participants to have completed the Ohio State University TBI Identification Screening Tool (OSU TBI-ID), along with PET amyloid imaging and APOE genotyping. Individuals were only excluded if they lacked any of the aforementioned three variables via missing data or failed quality control. As PET amyloid was the central focus for this project, included participants were not limited by tau PET availability. Informed consent was secured from all participants and received approval from the Human Research Protection Office at Washington University School of Medicine.

### Clinical Dementia Rating® (CDR®)

The cognitive status of participants was clinically evaluated using the Clinical Dementia Rating scale (CDR)®, which was administered by skilled clinicians through semi-structured interviews conducted with each participant and an informed collateral informant. The CDR was utilized to assess the extent of cognitive and functional impairment in participants[21]. A global CDR score of 0 signifies no cognitive impairment, a score of 0.5 reflects very mild dementia, while a score of 1 or higher indicates mild, moderate, or severe dementia.

### APOE Status

Extant APOE genotyping was acquired from blood samples gathered from all participants. In all participants, APOE was quantified according to the number of ε4 alleles (none, one, or two) and referring to APOE for the remainder of this article relates to the scaler 0-2 scale.

### Ohio State University TBI Identification Screening Tool

All participants underwent interviews conducted by trained research coordinators who strictly followed the specified guidelines to ensure uniform data collection across individuals[22]. All members of the research team responsible for collecting TBI-related data were blinded to the amyloid status or accumulation details. Participants were asked to provide details regarding any prior head injuries. Participants who reported a prior head injury were subsequently asked to provide details concerning the age at which the injury occurred and its severity, including the duration of any disoriented or fugue-like state and/or the length of time unconsciousness persisted. Due to limited mean distribution of the responses, the cohort was not further partitioned based on length of time associated with the fugue-like state or loss of consciousness. All incidents were documented and classification was determined based on the most recent and severe event. Individuals who did not report a TBI were categorized as “no TBI”, while those reporting a TBI were classified according to severity, with loss of consciousness (TBI-LOC) being considered more severe than experiencing a dazed state or fugue-like condition (TBI-DZ) following the incident[23]. The interval between the reported age of the most recent severe TBI and the date of the PET scan defined the recency of the TBI (greater than 10 or less than 10 years).

### Positron Emission Tomography

Aß PET was acquired with either 11C-Pitsuburgh Compound B (PiB)[24], or 18F-florbetapir (FBP)[25]. PiB was administered at a dose of approximately 13.94 mCi (SD=3.76), and analyzed from a 30-minute acquisition window 30 minutes post-injection. FBP was administered at a dose of approximately 9.92 mCi (SD=0.63), and analyzed from a 20-minute acquisition window 50 minutes after injection. All PET images were processed using the MRI-free PET processing pipeline by aligning the data to the MNI-152 atlas using the SPM12 coregister module[26], [27]. A global standard uptake value ratio (SUVr) was generated using the cortical ROI with the whole cerebellum as the reference region and then converting to the centiloid scale..

PET tau was acquired with 18F-Flortaucipir (FTP)[28] and analyzed from a 20-minute acquisition window 80 minutes post-injection. An SUVR global value was generated using the cerebellum-cortex as the reference region.

### White Matter Hyperintensities

All fluid attenuated inversion recovery (FLAIR) images were pre-processed using the following steps: brain extraction of the T1-weighted image, rigid body registration of the T1-weighted image to the corresponding FLAIR, and intensity normalization. WMH maps were then constructed with an 18-layer convolutional neural network with U-Net architecture. Further details on the full segmentation process have been described elsewhere[29].

## Statistics

### Primary Analyses

*The Recency of TBI on Amyloid-*β *Positivity;* A centiloid threshold exceeding 18.42 was established as the biomarker positivity criterion based on prior research findings[30]. Participants who reported a history of TBI were instructed to provide the age and severity of the injury, wherein loss of consciousness was identified as the primary characteristic determining severity followed by TBI-DZ or no TBI, referred to as the defining incident. For the analysis of TBI recency, individuals with TBI were categorized into two groups based on whether the classifying incident occurred within a decade prior to the PET scan or more than a decade before the PET scan. The cut-off of a single decade for defining remote vs recent was a-priori defined from prior work on this topic[31]. A Fisher’s exact test was conducted on the entire cohort and further stratified based on CDR status (CDR = 0 versus CDR > 0) to evaluate the association between amyloid positivity and the recency of TBI (<10 years versus >10 years). The analysis was performed on the entire TBI cohort and subsequently repeated for each distinct type of TBI individually.

#### Association Between

*TBI Severity and Amyloid:* Logistic and linear regression analyses were conducted to evaluate the relationship between TBI and amyloid positivity or centiloid levels, while adjusting for age, sex, education, and APOE ε4 alleles, stratified by CDR classification. The long-term effects of TBI were evaluated separately, incorporating distinct categorical variables for participants who indicated experiencing a fugue state without a loss of consciousness (TBI-DZ) and those who reported a loss of consciousness (TBI-LOC) resulting from the TBI incident. The primary regression model incorporated an interaction term to evaluate whether APOE ε4 amplifies the impact of chronic traumatic brain injury on amyloid positivity or accumulation. Pairwise post-hoc comparisons were carried out among TBI groups categorized based on CDR and APOE ε4. For the post-hoc comparisons, correction for multiple comparisons was adjusted using Bonferroni correction. All analyses were conducted using Matlab.

#### TBI Recency influences on APOE and TBI interaction

In the primary regression model for participants with a CDR= 0, analyses were further stratified based on TBI recency using the same criteria as described above. Using the same covariates and outcome measure, the regression analysis was conducted exclusively on the previously mentioned subsets to assess the strength of the association between AD biomarkers and timing of the TBI. Bonferroni correction was used to address multiple comparisons.

### Secondary Analyses

A visual graphic for changes in the sample size for any of the secondary analyses is provided in Supplemental Figure 1.

#### Spatial Specificity of TBI and AD

The amyloid centiloid mask was derived from well-characterized AD regions. However, to determine whether any potential relationship between TBI and amyloid deposition was specific to typical AD progression or represents non-specific amyloid accumulation, the bilateral insula—an area not typically implicated in early AD changes—was chosen as a control region. This secondary analysis utilized PiB SUVR values to circumvent potential complications associated with regional centiloid harmonization. Consequently, individuals with only AV-45 scans were excluded from this analysis. Employing the same statistical framework, we assessed the primary regression model by replacing the AD global centiloid outcome with the bilateral insula SUVR. The analysis was performed on the entire PiB subset and stratified based on CDR status.

#### TBI severity association with WMH

Since WMH generally deviates from a normal Gaussian distribution, the total WMH volume was log-transformed to mitigate the impact of skewness. The evaluation of WMH and TBI severity employed comparable statistical methods to those outlined earlier, with WMH replaced as the outcome variable while retaining the same predictors, including age, sex, education, APOE ε4, TBI-DZ, TBI-LOC, and the interaction between APOE ε4 and LOC.

#### Association Between TBI Severity and PET tau

Linear regression analyses were conducted to evaluate the relationship between TBI and PET tau, with adjustments for age, sex, education, and APOE ε4, stratified by CDR classification. The regression model incorporated an interaction term to evaluate whether a genetic predisposition to Alzheimer’s disease amplifies the impact of chronic TBI on PET tau accumulation.

## Results

### Study cohort

#### Cohort Demographics

A total of 429 individuals with a past history of TBI underwent a single PET amyloid scan, including 317 individuals scanned using PiB and 112 using AV45. Most participants (n=365) were identified as cognitively normal (CDR = 0), while 64 individuals (15%) were impaired (CDR > 0). The subsequent analyses will concentrate on the cognitively normal cohort (CDR=0) with the findings from the symptomatic cohort are provided in supplemental materials. The median age among all individuals was 67 years, with females comprising a disproportionately higher percentage at 60%. Among individuals with a CDR=0, 36% were genotyped as APOE positive, a proportion higher than that observed in the general population (Table 1).

**Table 1:** Demographic Table.

|  | Overall<br>(n=365) | Control<br>(n=246) | TBI-DZ<br>(n=56) | TBI-LOC<br>(n=63) |
| --- | --- | --- | --- | --- |
| Age | 66.5 | 66.4 | 66.0 | 66.9 |
| Sex (M/F) | 147/221 | 89/157 | 23/33 | 35/28 |
| Race<br>(Cau/AA/As/U) | 330/36/3/1 | 219/27/2/0 | 50/5/0/1 | 58/4/1/0 |
| CDR-SoB<br>Average(SD) | 0.005 (0.05) | 0.008 (0.06) | 0 (0) | 0 (0) |
| Years of Education | 16 | 16 | 16 | 16 |
| APOE | 36% | 35% | 39% | 40% |
| TBI 1-10 yrs |  |  | 15 | 19 |
| TBI > 10 yrs |  |  | 41 | 44 |
| TBI Recency |  |  | 36.3 | 31.5 |

#### TBI Demographics

Individuals who reported a head injury without any alteration in cognitive state (i.e., dizziness [DZ] or loss of consciousness [LOC]) were classified as equivalent to those who reported no TBI and were collectively designated as the control group (Con). Among the 365 participants, 119 reported a history of TBI, while 246 did not report a TBI prior to undergoing PET imaging. A significant proportion of individuals with a TBI reported experiencing an LOC, either accompanied by or independent of dizziness (53%). No significant differences were identified between the asymptomatic chronic TBI group and Con with respect to age, race, education, or APOE status; however, the TBI-LOC group exhibited a higher proportion of males. No significant differences were observed in TBI recency within the TBI group (Table 1).

### Primary Analyses

#### The Recency of TBI on Amyloid-β Positivity

Within the Con group, 29 individuals (12%) were identified as amyloid positive, while 217 individuals (88%) were classified as amyloid negative. Among individuals in the TBI cohort, 33 participants reported sustaining a TBI within ten years prior to undergoing the PET amyloid scan. Of these, 8 (24%) exhibited pathological amyloid levels, while 25 (76%) demonstrated sub-pathological levels. Within the TBI group, most participants (n=85) reported a history of remote TBI, with 24 individuals (28%) testing positive for amyloid and 61 individuals (72%) testing negative for amyloid. An analysis of the timing of all TBIs, compared to the Con group, indicated that remote TBI was significantly correlated with increased amyloid accumulation (p<0.001). Conversely, recent TBI did not demonstrate statistical significance after adjusting for multiple comparison. When stratified by TBI severity, the analysis revealed that both recent and remote TBI-LOC were associated with amyloid accumulation (recent: p = 0.004; remote: p < 0.001) (Supplemental Table 1). However, neither association was statistically significant among individuals with TBI-DZ (recent: p = 0.1; remote: p = 0.29) (Supplemental Table 2).

#### Association Between TBI Severity and Amyloid positivity

Age and the presence of the APOE ε4 were found to be significantly associated with amyloid positivity (both p<0.001), whereas neither sex nor educational level demonstrated a significant contribution to the model. Previous exposure to a TBI-LOC is strongly correlated with amyloid positivity, exhibiting a comparable odds ratio to that of APOE ε4 (Figure 1A). Prior exposure to a TBI-DZ was not significantly associated with amyloid positivity, suggesting that not all TBI events are comparable. This correlation remained consistent when analyzed exclusively among asymptomatic individuals (CDR=0). Subsequent pairwise comparisons among the groups revealed that, within the full cohort, the TBI-LOC group exhibited a significantly higher proportion of amyloid-positive individuals (p < 0.001). However, this difference was not significant in the TBI-DZ group after correcting for multiple comparisons. Conversely, in individuals with CDR=0, the TBI-LOC group exhibited a significant difference when compared to both groups (p<0.001 vs Con; p=0.004 vs TBI-DZ). No significant findings were observed for the symptomatic cohort (CDR>0) and findings are provided in the supplementary materials (Supplemental Table 3).

**Figure 1:**
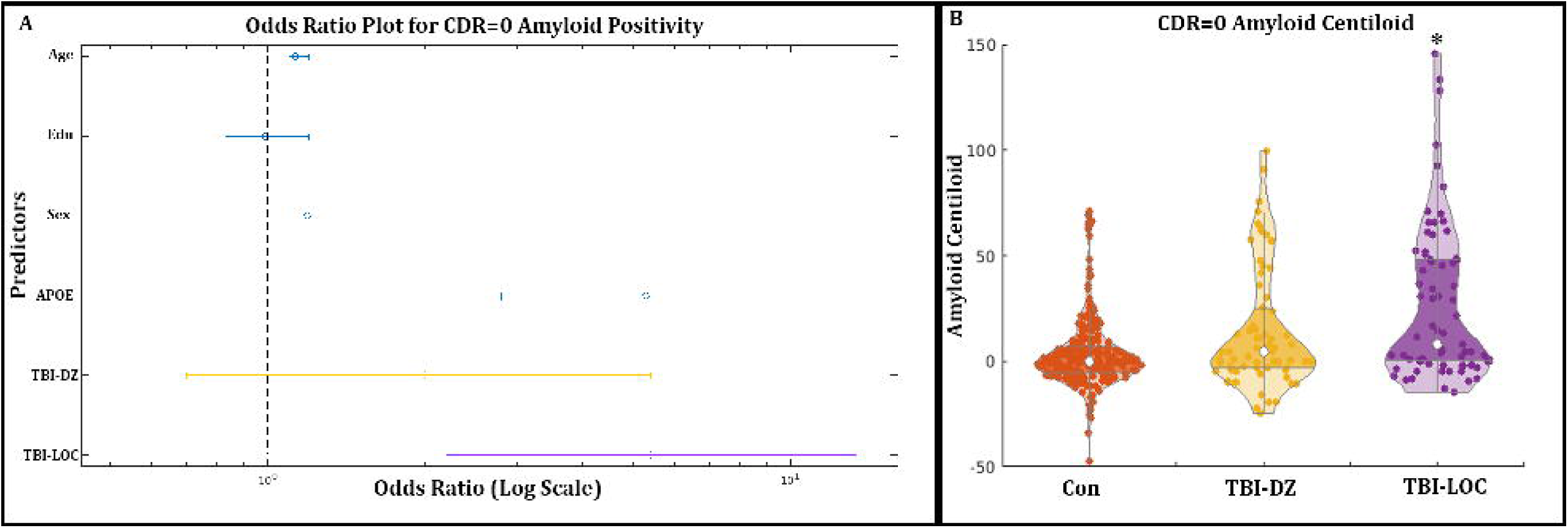
A) The significant predictors in the logistic regression for amyloid positivity were age, ApoE4 and LOC as demonstrated in the Odds Ratio plot. B) The elevated amyloid in LOC is visually displayed in Figure 1B with significantly higher amyloid in the LOC group compared to both Con (p<0.001) and TBI-DZ (p=0.004). *= Significantly different; TBI-DZ= Traumatic Brain injury with dazed or confusion; TBI-LOC = Traumatic Brain injury with Loss of Consciousness; Con = Controls; APOE = Apolipoprotein E4; CDR = Clinical Demetia Rating

**Figure 2:**
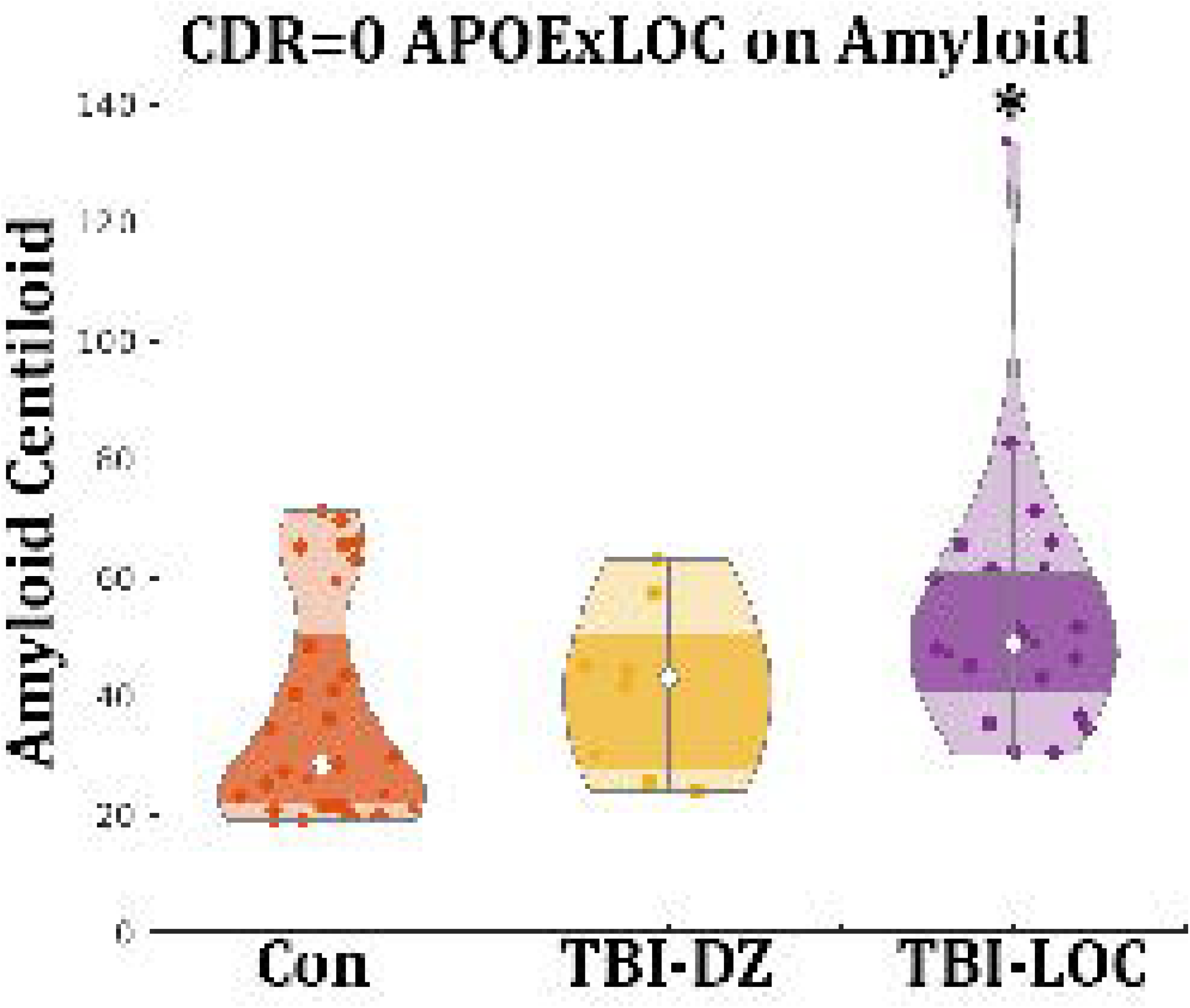
The interaction between ApoE4 and LOC is visually displayed where each data point represents an individual with at least one E4 allele. The TBI-LOC group was significantly different from the Con group (p=0.003) but not the TBI-DZ group.*= Significantly different; TBI-DZ= Traumatic Brain injury with dazed or confusion; TBI-LOC = Traumatic Brain injury with Loss of Consciousness; Con = Controls; APOE = Apolipoprotein E4; CDR = Clinical Demetia Rating

#### Association Between TBI Severity and Continuous Amyloid Levels

Analyzing amyloid levels as a continuous variable, following centiloid conversion within a linear model, demonstrated relationships comparable to those identified with biomarker positivity. Age and the presence of the APOE ε4 allele demonstrated a significant impact on amyloid accumulation (p<0.001), whereas sex did not exhibit a statistically significant effect. Education demonstrated a slight negative correlation with amyloid accumulation (p=0.033). A previous history of TBI-LOC was significantly associated with amyloid accumulation (p<0.001); however, no correlation was observed with TBI-DZ (p=0.62). Furthermore, no interaction between APOE ε4 and TBI-LOC was detected in the analysis of the entire cohort. In CDR=0 individuals age and the presence of the APOE ε4 persisted as significant predictors of amyloid accumulation (both p<0.001). The specificity of TBI remained evident, as only TBI-LOC was associated with amyloid accumulation (p<0.001) (Figure 1B). Moreover, a notable interaction between APOE ε4 and TBI-LOC was identified (p = 0.003) (Table 2). A single individual was identified as a potential outlier; however, the results remained statistically significant even after reanalyzing the data with their exclusion (Supplemental Table 4). No significant findings were observed for the symptomatic cohort (CDR>0) and findings are provided in the supplementary materials (Supplemental Table 4).

**Table 2:** Linear regression results for associating with Amyloid centiloids in CDR=0 participants.

| Predictors | Estimates | CI | tstat | p |
| --- | --- | --- | --- | --- |
| (Intercept) | -6.3 | -9.3-(-3.3) | -4.1 | **<0.001** |
| age | 0.50 | 0.31-0.72 | 5.0 | **<0.001** |
| Sex | 0.4 | -3.5-4.3 | 0.2 | 0.88 |
| Edu | 0.39 | -0.46-1.2 | 0.9 | 0.28 |
| Allele (APOE) | 7.6 | 4.0-11 | 4.1 | **<0.001** |
| DZ | 1.7 | -3.6-6.9 | 0.63 | 0.53 |
| LOC | 10.7 | 4.5-17 | 3.4 | **<0.001** |
| APOE:LOC | 12.7 | 4.2-21 | 2.9 | **0.003** |

### TBI Recency influences on APOE Genotype and TBI interaction

A confined analysis to participants with either recent or remote TBI-LOC indicated a significant main effect for recent TBI-LOC (p = 0.015), APOE ε4 (p < 0.001), and age (p < 0.001). However, the interaction between APOE ε4 and LOC was not statistically significant (p = 0.67) (Supplemental table 6). Conversely, while the same factors were significant for remote TBI-LOC, a notable interaction was observed between APOE ε4 and LOC (p=0.003) (Supplemental 7).

### Secondary Analyses

#### Spatial Specificity of TBI and AD

The spatial specificity for AD was assessed using the PiB SUVR for the bilateral Insula as a non-AD specific early-stage control region. No association was observed between amyloid accumulation and TBI severity for either the full cohort or the CDR=0 subset.

#### TBI severity association with WMH

The integrity of white matter was evaluated using global WMH as a proxy, employing the same predictor variables outlined in the primary linear regression model. Significant main effects were identified solely for age (p<0.001) and APOE ε4 (p=0.035), while no such effects were observed for either variant of TBI. Nevertheless, a notable interaction between APOE ε4 and LOC was identified (p=0.013), suggesting a synergistic influence on the degeneration of white matter (Table 3).

**Table 3:** Linear regression results for associating with white matter hyperintensities in CDR=0 participants.

| Predictors | Estimates | CI | tstat | p |
| --- | --- | --- | --- | --- |
| (Intercept) | 0.013 | -0.12-0.14 | 0.21 | 0.84 |
| age | 0.050 | 0.041-0.058 | 11 | <**0.001** |
| Sex | 0.05 | -0.11-0.22 | 0.66 | 0.51 |
| Edu | -0.013 | -0.05-0.02 | -0.69 | 0.49 |
| Allele (APOE) | -0.17 | -0.33-(-0.012) | -2.1 | **0.035** |
| DZ | 0.17 | -0.05-0.39 | 1.49 | 0.14 |
| LOC | -0.17 | -0.43-0.10 | -1.2 | 0.22 |
| APOE:LOC | 0.46 | 0.099-0.82 | 2.5 | **0.013** |

#### TBI Severity and Tau PET

Substituting the outcome measure with PET tau SUVR values did not reveal any significant association with either of the TBI groups. This absence of statistical significance was identified in both the entire cohort and the CDR=0 subgroup. Findings from the association with the tau summary metric can be found in supplemental table 8.

## Discussion

This extensive evaluation of TBI and AD biomarker revealed that cognitively unimpaired individuals with a history of remote LOC who carry at least one APOE ε4 allele are at risk of increased amyloid deposition,. Notably, these findings were not observed among symptomatic individuals (CDR > 0). Moreover, this relationship exhibited spatial specificity for AD-like amyloid deposition, as no significant association was identified in the non-AD control region. Although few studies have provided evidence of elevated neurofibrillary tangles in participants with history of TBI[32][33], we did not detect any association with tau PET for any Braak stage, either across the entire cohort or when stratified by CDR status. However, elevated WMH presence was observed in individuals with TBI-LOC and APOE. Collectively, these findings suggest a specific amyloid-centered mechanism characterized by a gene-environment “dual-hit,” involving TBI-LOC and the APOE ε4 allele.

The link between amyloid deposition and TBI remains unclear but has emerged in various cohorts with different experimental designs. In post-mortem studies, amyloid plaques have been detected following recent injuries with increasing deposition depending on the age at death[3], [34]. Moreover, postmortem analyses of individuals who survived TBI over the long term have revealed the existence of both amyloid-beta plaques and neurofibrillary tangles[33]. In vivo studies of acute traumatic brain injury (TBI) have also identified increased amyloid deposition, frequently occurring within days post-injury[4]. In the case of chronic TBI observation of AD biomarkers reveal inconsistent evidence regarding amyloid deposition in cases of chronic TBI a decade post-incident[5]. We propose that the observed inconsistency arises from the genetic impact of APOE ε4, which, in this study, was synergistically linked with increased amyloid levels in conjunction with TBI.

The synergistic interaction between TBI and APOE ε4 might be explained by hypotheses concerning axonal injury, with previous studies indicating that APOE ε4 has an influential role in restoration of axonal integrity following acute TBI[35]. Research utilizing animal models of acute TBI indicates that APOE ε4 contributes to the reparative process following a TBI[35]. However, the efficacy of recovery appears to vary depending on the genotype, with APOE ε4 being associated with longer recovery response times compared to APOE ε3, ultimately leading to a less favorable prognosis [36]. Moreover, studies involving humans and suitable animal models have demonstrated that the APOE e4 allele is associated with dysfunctional oligodendrocytes and consequently defective myelin sheaths surrounding the axons[37]. Research on aging has demonstrated that individuals with the APOE gene may experience a decline in the structural integrity of myelin sheaths during midlife, which is exacerbated with multiple copies of the e4 allele and subsequently contribute to the development of neurodegenerative disorders in later stages of life [38], [39], [40]. Additionally, recovery from TBI has been observed to pose greater challenges in pediatric APOE ε4 patients compared to their peers of similar age without this genotype, indicating a potential lifelong vulnerability to TBI[39]. We observed a significant coupling of TBI and APOE ε4 for WMH frequency that was used as a proxy for white matter integrity but future studies incorporating diffusion tensor models are needed to validate this finding.

The relationship between APOE ε4 and TBI has been demonstrated to vary based on the outcome variable, with biomarker outcome measures being more prone to detect a significant association[41]. This aligns with our findings, which demonstrated that this relationship was evident solely among cognitively normal individuals and absent in those who were symptomatic. While similar findings were observed across the entire cohort, the relationship remained evident only among cognitively normal individuals when analyzed based on CDR status. Similarly, prior work in the Alzheimer Disease Neuroimaging Initiative (ADNI) reported elevated amyloid for cognitively normal TBI compared to controls but no difference was observed between individuals with or without TBI in CDR>0 individuals[42].

As AD advances, the accumulation of amyloid plateaus, while neurofibrillary tangles emerge as the predominant pathological feature, exhibiting a stronger association with cognitive decline and neurodegenerative processes. However, the associations observed with PET tau remain inconsistent, as some studies report modest correlations, while no associations are identified in well-defined civilian[43], [44] and military cohorts[16]. This study identified no correlation between PET tau and TBI, suggesting that the mechanisms associated with TBI may have a reduced impact during the advanced stages of the disease. Although our findings were predominantly observed in asymptomatic individuals, where neurofibrillary tangles (NFTs) are generally known to be at relatively low levels, no association was identified within the symptomatic cohort either. Alternatively, regardless of the origin of amyloid accumulation, once individuals reach a certain stage, the progression of Alzheimer’s disease becomes indistinguishable from mechanisms induced by TBI and other variants of sporadic AD. Nonetheless, additional longitudinal research is required to substantiate these assertions. TBI is typically regarded as associated to an increased likelihood to all dementia however, our findings revealed a disease-specific spatial correlation with amyloid accumulation that was absent in the non-AD control region. This is congruent with other studies that has evaluated the spatial distribution of AB deposition in TBI individuals [45] but atypical increases in the cerebellum has also been reported[4]. We selected the insula as our control region since the cerebellum was used as the reference region for the SUVR calculations.

Several limitations exist that warrant attention, as future research designed to assess these claims would be beneficial. This cohort exhibited an increased prevalence of AD, with a higher frequency of the APOE ε4 allele compared to that observed in the general population. Replicating these findings within a larger cohort would significantly strengthen the evidence for the interaction between TBI-LOC and APOE ε4. This study does not include adequate longitudinal follow-up to ascertain whether individuals with a history of TBI-LOC and genetic predisposition to AD exhibit a higher incidence rate or faster progression to AD compared to their respective control groups. TBI history was acquired through self-report and not verified with medical records which is subject to scrutiny and more dedicated cohorts to chronic TBI are needed to validate these claims. Additionally, inflammatory measures were not included in this study which are of growing interest mechanistically between TBI and AD. A longitudinal study could yield more comprehensive insights into whether the observed increase in amyloid levels among these individuals is associated with clinically significant changes over time. Although no association with tau was observed our sample had considerably more cognitively normal individuals which can influence that outcome regarding the synergy between tau and cognitive decline. The history of TBI was obtained through self-reporting, which is less reliable than medical records and prone to variability due to the absence of a verified source or adequate documentation. Efforts were made to mitigate potential bias by refraining from assessing the frequency of TBI and instead concentrating on singular events that were associated with changes in cognitive state (DZ and LOC). Previous research in this field indicates that while self-reporting possesses certain limitations, the accuracy of self-reports regarding the experience of ever having suffered a TBI-LOC is comparatively more reliable. Research with more precise documentation would facilitate the opportunity to conduct more comprehensive analyses of the specifics of these injuries.

This study indicates that remote TBI-LOC is correlated with increased amyloid PET levels in later life, particularly among APOE ε4 carriers with a CDR of 0, thereby supporting a synergistic “dual-hit” hypothesis where TBI-LOC exacerbates APOE ε4-driven susceptibility to amyloid accumulation. The selectivity for amyloid (as opposed to tau) and its diminished presence in individuals with CDR>0 underscore a sustained, amyloid-centered mechanism. In research and clinical contexts, PET amyloid positivity continues to be the benchmark for defining preclinical AD cohorts. Furthermore, while positivity is a requirement for eligibility in anti-amyloid therapy, a history of TBI is not currently taken into account. The impact of APOE is well-established and evident in our study, showing a general rise in amyloid accumulation among all individuals, irrespective of their TBI history. Nevertheless, when paired with TBI-LOC, the influence of APOE ε4 on amyloid accumulation became more pronounced, even in cases where the TBI occurred more than a decade earlier. This underscores the importance of obtaining and factoring in TBI information when assessing or identifying preclinical AD cohorts for clinical relevance.

## Acknowledgements

This study was primarily funded through the Congressionally directed Medical Research Programs (CDMRP) as a subsidiary of the DoD (AZ220074 and AZ230119). A portion of the project was funded in part by the National Institutes of Health (NIH; P01-AG026276, P01-AG03991, P30-AGG066444, K01-AG090753 [PI: PRM]). It is subject to the NIH Public Access Policy. Through acceptance of this federal funding, NIH has been given a right to make this manuscript publicly available in PubMed Central upon the Official Date of Publication, as defined by NIH. This work was only possible due to the participants and the personnel of the Knight ADRC for providing this comprehensive dataset used in this project.

