## Supplemental for "History of Traumatic Brain Injury with Loss of Consciousness and APOE ε4 Carriers Synergistically Increase Late-Life Amyloid PET Burden"

Sample Size Breakdown by Analysis

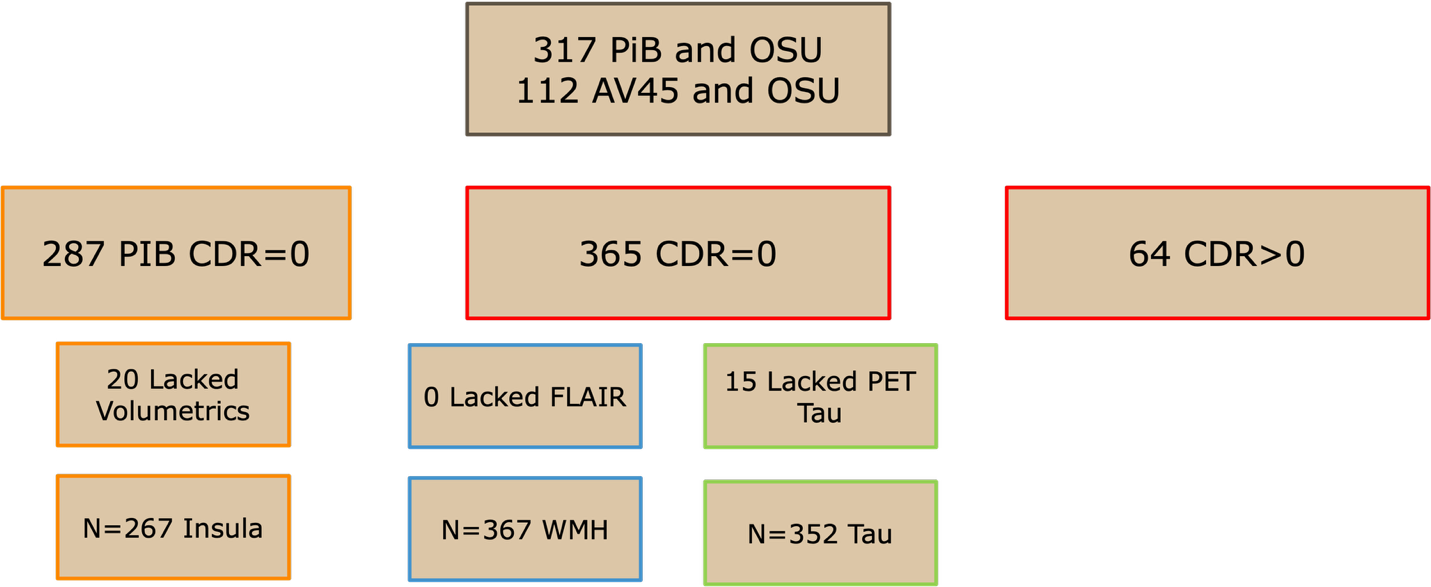

Supplemental Figure 1: Cohort breakdown by analysis. Boxes highlighted in red represent the sample sizes used in the primary analyses of amyloid centiloid/positivity. Blue boxes represent the sample size for the WMH analysis and the green boxes represent the sample size for the PET tau analysis. The AD spatial analysis was only performed on the PiB individuals, which is represented by the boxes with an orange outline. PiB = Pittsburgh compound B; OSU = Ohio State University TBI screening Tool; AV45= florbetapir; CDR = Clinical Dementio Ratio; FLAIR = Fluid attenuated inversion recovery scan; WMH = white matter hyperintensity; PET = Positron Emission Tomography.

| TBI-LOC | **Aβ +**  **(n=53)** | **Aβ –**  **(n=287)** |
| --- | --- | --- |
| **TBI History** | | |
| TBI 1-10 years | 7 (37%) | 12 (63%) |
| **TBI Past 10 Years** | **P=0.004** | |
| TBI > 10 years | 17 (39%) | 27 (61%) |
| **TBI >10 Years** | **P<0.001** | |
| Con | 29 (10%) | 248 (90%) |

Supplemental Table 1: Fisher’s exact test on amyloid positivity in TBI-LOC compared to the Con group confined to CDR=0 participants and stratified by TBI Recency. The top half shows the frequency any positivity percentage of recent TBI-LOC and the bottom half shows the frequency and positivity percentage of remote TBI-LOC. TBI-LOC = Traumatic Brain injury with Loss of Consciousness; Con = Controls; Aβ = amyloid-beta

| TBI-DZ | **Aβ +**  **(n=37)** | **Aβ –**  **(n=296)** |
| --- | --- | --- |
| **TBI History** | | |
| TBI 1-10 years | 1 (0.07%) | 14 (93%) |
| **TBI Past 10 Years** | P=0.1 | |
| TBI > 10 years | 7 (17%) | 34 (83%) |
| **TBI >10 Years** | P=0.29 | |
| Con | 29 (10%) | 248 (90%) |

Supplemental Table 2: Fisher’s exact test on amyloid positivity in TBI-DZ compared to the Con group confined to CDR=0 participants and stratified by TBI Recency. The top half shows the frequency any positivity percentage of recent TBI-DZ and the bottom half shows the frequency and positivity percentage of remote TBI-DZ. TBI-DZ= Traumatic Brain injury with dazed or confusion; Con = Controls; Aβ = amyloid-beta

| **Predictors** | **Odds Ratios** | **CI** | **p** |
| --- | --- | --- | --- |
| (Intercept) | 0.015 | 6.7e-9-34869 | 0.56 |
| Age | 1.11 | 0.93-1.3 | 0.22 |
| Allele (APOE) | 1.6 | 0.30-9.1 | 0.55 |
| Edu | 0.94 | 0.59-1.5 | 0.78 |
| Sex | 0.28 | 0.019-4.0 | 0.32 |
| TBI-LOC | 1.5 | 0.10-24.6 | 0.75 |
| TBI-DZ | 0.15 | 0.0062-3.5 | 0.21 |

Supplemental Table 3: Logistic Regression on amyloid positivity for CDR>0 participants. TBI-DZ= Traumatic Brain injury with dazed or confusion; TBI-LOC = Traumatic Brain injury with Loss of Consciousness; Con = Controls; APOE = Apolipoprotein E4; CDR = Clinical Demetia Rating; Edu = education

| **Predictors** | **Estimates** | **CI** | **tstat** | **p** |
| --- | --- | --- | --- | --- |
| (Intercept) | -5.6 | -8.5-(-2.8) | -3.8 | **<0.001** |
| age | 0.45 | 0.26-0.65 | 4.6 | **<0.001** |
| Sex | -0.16 | -3.8-3.5 | -0.087 | 0.93 |
| Edu | 0.77 | -0.04-1.6 | 1.9 | 0.063 |
| Allele (APOE) | 7.4 | 3.9-11 | 4.1 | **<0.001** |
| DZ | 1.8 | -3.1-6.8 | 0.72 | 0.47 |
| LOC | 10 | 4.3-16 | 3.4 | **<0.001** |
| APOE:LOC | 10 | 2.1-18 | 2.5 | **0.01** |

Supplemental Table 4: Linear Regression on amyloid centiloids with the potential outlier removed in CDR=0 participants. TBI-DZ= Traumatic Brain injury with dazed or confusion; TBI-LOC = Traumatic Brain injury with Loss of Consciousness; Con = Controls; APOE = Apolipoprotein E4; CDR = Clinical Demetia Rating; Edu = education

| **Predictors** | **Estimates** | **CI** | **tstat** | **p** |
| --- | --- | --- | --- | --- |
| (Intercept) | 99 | -66-264 | 1.2 | 0.23 |
| Age | -0.37 | -2.3-1.6 | -0.38 | 0.7 |
| Sex | 4.8 | -20.3-30 | 0.38 | 0.7 |
| Edu | -2.4 | -6.8-2.0 | -1.1 | 0.28 |
| Allele (APOE) | 23 | 1.7-43 | 2.2 | **0.034** |
| TBI-DZ | 7.5 | -25-40 | 0.46 | 0.65 |
| TBI-LOC | 19 | -25-63 | 0.87 | 0.39 |
| APOE:LOC | -25 | -65-15 | -1.3 | 0.21 |

Supplemental Table 5: Linear Regression on amyloid centiloid for CDR>0 participants. TBI-DZ= Traumatic Brain injury with dazed or confusion; TBI-LOC = Traumatic Brain injury with Loss of Consciousness; Con = Controls; APOE = Apolipoprotein E4; CDR = Clinical Demetia Rating; Edu = education

| **Predictors** | **Estimates** | **CI** | **tstat** | **p** |
| --- | --- | --- | --- | --- |
| (Intercept) | -5.9 | -8.9-(-2.8) | -3.8 | **<0.001** |
| age | 0.52 | 0.31-0.73 | 4.8 | **<0.001** |
| Sex | 2.8 | -1.1-6.8 | 1.4 | 0.16 |
| Edu | 0.56 | 0.32-1.4 | 1.3 | 0.21 |
| Allele (APOE) | 10 | 6.6-14 | 5.6 | **<0.001** |
| Recent TBI-DZ | -0.78 | -6.2-4.6 | -0.28 | -0.78 |
| Recent TBI-LOC | 13 | 2.6-24 | 2.4 | **0.015** |
| APOE:LOC | 3.2 | -12-18 | -0.43 | 0.67 |

Supplemental Table 6: Linear Regression on amyloid centiloids in CDR=0 participants. Recent TBI-DZ= Traumatic Brain injury with dazed or confusion that occurred within 10 years from the PET scan; Recent TBI-LOC = Traumatic Brain injury with Loss of Consciousness that occurred within 10 years from the PET scan; Con = Controls; APOE = Apolipoprotein E4; CDR = Clinical Demetia Rating; Edu = education

| **Predictors** | **Estimates** | **CI** | **tstat** | **p** |
| --- | --- | --- | --- | --- |
| (Intercept) | -5.1 | -8.1-(-2.1) | -3.4 | **<0.001** |
| age | 0.49 | 0.28-0.70 | 4.7 | **<0.001** |
| Sex | 0.022 | -3.9-4.0 | 0.011 | 0.99 |
| Edu | -0.61 | -0.25-1.5 | 1.4 | 0.16 |
| Allele (APOE) | 8.0 | 4.4-12 | 4.4 | **<0.001** |
| Remote TBI-DZ | 0.54 | -4.8 | 0.20 | 0.84 |
| Remote TBI-LOC | 8.1 | 0.64-15 | 2.1 | **0.033** |
| APOE:LOC | 15 | 5.2-25 | 3.0 | **0.003** |

Supplemental Table 7: Linear Regression on amyloid centiloids in CDR=0 participants. Remote TBI-DZ= Traumatic Brain injury with dazed or confusion that occurred more than 10 years from the PET scan; Remote TBI-LOC = Traumatic Brain injury with Loss of Consciousness that occurred more than 10 years from the PET scan; Con = Controls; APOE = Apolipoprotein E4; CDR = Clinical Demetia Rating; Edu = education

| **Predictors** | **Estimates** | **CI** | **tstat** | **p** |
| --- | --- | --- | --- | --- |
| (Intercept) | 0.027 | -0.005-0.058 | 1.7 | 0.09 |
| age | 0.0048 | 0.0027-0.0069 | 4.4 | **<0.001** |
| Sex | -0.1 | -0.14-(-0.057) | -4.7 | **<0.001** |
| Edu | 0.002 | -0.0064-0.011 | 0.55 | 0.58 |
| Allele (APOE) | 0.035 | 0.0044-0.073 | 1.7 | 0.082 |
| TBI-DZ | -0.03 | -0.084-0.024 | -1.1 | 0.27 |
| TBI-LOC | -0.01 | -0.075-0.052 | -0.35 | 0.73 |
| APOE:LOC | 0.06 | -0.022-0.15 | 1.5 | 0.14 |

Supplemental Table 8: Linear Regression on PET tau in Braak Stage 1-2 for all participants. TBI-DZ= Traumatic Brain injury with dazed or confusion; TBI-LOC = Traumatic Brain injury with Loss of Consciousness; Con = Controls; APOE = Apolipoprotein E4; CDR = Clinical Demetia Rating; Edu = education
